# MOSurvivor-Guided Joint CpG Selection and XGBoost Hyperparameter Optimization for Compact Epigenetic Age Prediction

**DOI:** 10.64898/2026.08.26.747213

**Authors:** Arif Yelği, Shirmohammad Tavangari, Zahra Shakarami, Sajjad Janfaza

**Author notes:** Corresponding author: Arif Yelği.

## Abstract

Accurate epigenetic age prediction from DNA methylation profiles is intrinsically high-dimensional, creating a need for parsimonious models that preserve predictive performance while reducing the number of assayed cytosine-phosphate-guanine (CpG) loci. This study introduces MOSurvivor, a population-based multi-objective search framework that jointly optimizes a weight-threshold CpG selector and eight XGBoost hyperparameters. Experiments used the GSE40279 whole-blood cohort (656 individuals profiled on the Illumina HumanMethylation450 platform). After retaining 1,000 age-correlated CpGs, five strategies were evaluated on the same 30 seeded 80:20 train/test splits: fixed-parameter XGBoost using all 1,000 CpGs, random search, a genetic algorithm, particle swarm optimization, and MOSurvivor. Internal fitness was estimated using three-fold cross-validation on each training set. Across the 30 held-out test sets, MOSurvivor achieved a mean absolute error (MAE) of 4.149 ± 0.300 years, root mean squared error of 5.545 ± 0.392 years, and R^2^ of 0.855 ± 0.027 while retaining 211.6 ± 54.8 CpGs. Relative to full-feature XGBoost (MAE 4.095 ± 0.285 years), MOSurvivor reduced the feature set by 78.8% at an MAE increase of only 0.054 years (1.3%). Paired Wilcoxon tests found no significant accuracy difference between MOSurvivor and any comparator (all unadjusted p > 0.05; all Holm-adjusted p ≥ 0.476). The most recurrent locus, cg16867657, appeared in 29 runs, whereas mean pairwise Jaccard similarity was 0.124, indicating a small stable core embedded in multiple near-equivalent feature subsets. MOSurvivor thus offers a competitive accuracy-parsimony trade-off rather than superior absolute accuracy. External validation and leakage-free nested feature preselection remain necessary before biological or clinical translation.

## 1. Introduction

DNA methylation (DNAm) changes reproducibly with age and can be summarized into epigenetic clocks that estimate chronological or biological age. Early whole-blood and multi-tissue clocks demonstrated that a comparatively small set of CpG loci can capture a substantial portion of age-related variation [1–3]. These models have become important tools in aging research, epidemiology, and studies of age acceleration, but their construction remains challenging because methylation arrays measure hundreds of thousands of candidate CpGs in cohorts that are usually orders of magnitude smaller.

Penalized linear models are widely used for epigenetic clocks because they perform embedded feature selection, yet non-linear learners may capture interactions and threshold effects that are missed by strictly additive models. XGBoost provides a flexible and computationally efficient tree-boosting framework [4], but its performance depends on both the selected features and the hyperparameter configuration. Optimizing these two components sequentially can be suboptimal: the best hyperparameters for a broad CpG panel need not be optimal for a compact panel, and vice versa.

Population-based metaheuristics offer a natural mechanism for mixed-variable joint optimization. However, using such algorithms solely to minimize prediction error can yield unnecessarily large feature subsets, while aggressive sparsity can compromise accuracy. The present work evaluates MOSurvivor, a multi-objective Survivor-inspired optimizer that encodes CpG weights, an adaptive selection threshold, and XGBoost hyperparameters in a single candidate. The principal hypothesis is not that MOSurvivor must outperform a full-feature model in absolute accuracy, but that it can preserve statistically comparable test performance with substantially fewer CpGs.

The contributions are fourfold: (i) a unified mixed representation for CpG selection and XGBoost tuning; (ii) an adaptive threshold and repair mechanism that constrains subset size; (iii) a 30-run paired comparison against full-feature XGBoost and three search baselines; and (iv) a stability analysis of recurrent CpGs and pairwise overlap across independent data splits.

## 2. Materials and Methods

### 2.1 Dataset and prediction target

GSE40279 contains genome-wide DNAm profiles from whole blood of 656 human individuals spanning a broad adult age range, measured with the Illumina Infinium HumanMethylation450 BeadChip [1,5]. The response variable was chronological age in years, and CpG beta values served as predictors. The public dataset contains no intervention; the present analysis is a secondary computational study of de-identified data.

### 2.2 Preprocessing and experimental design

The 1,000 CpGs with the largest absolute correlation with age were retained as an initial screening panel. For each of 30 independent runs, a seed from 42 through 71 generated an 80% training and 20% test split. The same split in each run was shared across all five methods, enabling paired comparisons. Candidate solutions were scored using three-fold cross-validation within the training partition, after which the selected configuration was refitted on the complete training partition and evaluated once on its held-out test set.

One validity concern deserves emphasis: in the supplied pipeline, the initial correlation screen was performed before the train/test split. Because age information from the complete dataset contributed to this screen, the reported errors may be optimistic. This issue does not invalidate the within-experiment paired comparison, since every method received the same 1,000-CpG panel, but it limits generalization claims. A confirmatory study should repeat the screen independently inside each training split or within an outer nested-validation loop.

### 2.3 Candidate representation and XGBoost search space

Each MOSurvivor candidate comprised three blocks: 1,000 continuous CpG weights, a continuous selection threshold, and eight XGBoost hyperparameters. A CpG was activated when its weight exceeded the candidate-specific threshold. The model block encoded maximum tree depth, learning rate, number of estimators, subsample ratio, column-sampling ratio, minimum child weight, gamma, and L2 regularization (reg_lambda). A repair operator enforced 50–300 selected CpGs, preventing degenerate or excessively dense candidates.

**Table 1.** Mixed representation used by MOSurvivor.

| Component | Decision variables | Role |
| --- | --- | --- |
| CpG selector | 1,000 continuous weights | Scores candidate CpG loci |
| Adaptive threshold | 1 continuous value | Converts weights to a binary subset |
| XGBoost model | 8 hyperparameters | Controls tree complexity, shrinkage, sampling, and regularization |
| Repair constraint | 50–300 CpGs | Maintains feasible subset size |

### 2.4 MOSurvivor optimization

The population was divided into two groups. Pairwise competitions identified winners and losers, and candidate updates learned from the pairwise winner, the best member of the local group, the best global member, and leaders sampled from a Pareto archive. The stochastic perturbation rate decreased over iterations, shifting the search from exploration toward exploitation. Predictive error was prioritized while subset size acted as a secondary parsimony objective. For final solution selection, the scalar score was MAE_CV + 0.10 × (number of selected CpGs / 1,000); thus, one additional year of cross-validated MAE carried substantially more weight than moderate differences in subset size.

### 2.5 Comparator methods

MOSurvivor was compared with: (i) a fixed-parameter XGBoost model fitted to all 1,000 screened CpGs; (ii) random search over the joint weight-threshold and hyperparameter representation; (iii) a genetic algorithm (GA); and (iv) particle swarm optimization (PSO). The three optimized baselines performed both subset selection and XGBoost tuning. The available output files do not document an exactly equalized number of model evaluations across optimizers; computational-efficiency claims are therefore intentionally avoided.

### 2.6 Evaluation and statistical analysis

Held-out prediction was summarized using MAE, root mean squared error (RMSE), and the coefficient of determination (R^2^). Feature parsimony was summarized as the number of selected CpGs and percentage reduction relative to 1,000 CpGs. Because all methods were evaluated on matched splits, two-sided paired Wilcoxon signed-rank tests compared MOSurvivor with each comparator. Holm correction controlled the family-wise error rate across four comparisons. Stability was described using selection frequency and the Jaccard index |A ∩ B| / |A ∪ B| for every pair among the 30 MOSurvivor subsets.

## 3. Results

### 3.1 Predictive performance and parsimony

The full-feature XGBoost model attained the lowest mean MAE (4.095 years), followed by PSO (4.124), MOSurvivor (4.149), GA (4.170), and random search (4.197). MOSurvivor used the smallest mean subset among the optimized methods: 211.6 CpGs versus 293.4 for PSO, 269.6 for GA, and 278.7 for random search. Relative to the 1,000-CpG reference, this corresponds to a 78.8% reduction in dimensionality for an absolute MAE increase of 0.054 years (approximately 19.8 days) and a relative increase of 1.3%.

**Table 2.**
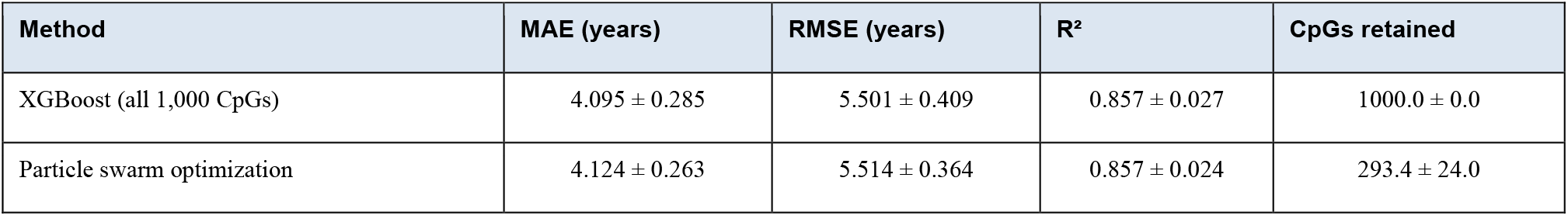
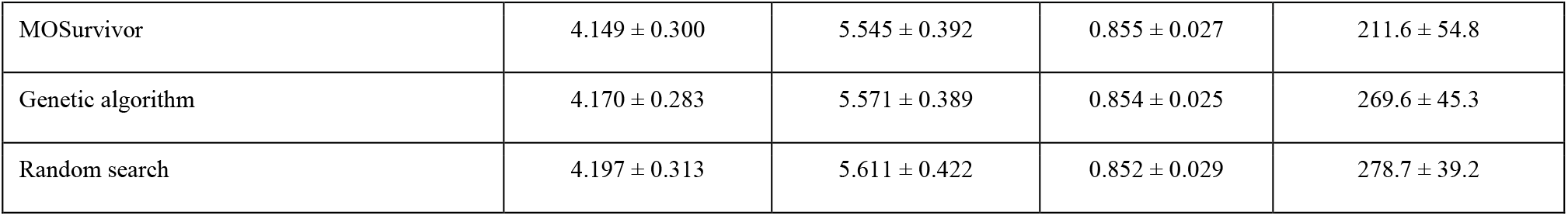
Held-out performance over 30 matched train/test splits (mean ± SD).

| Method | MAE (years) | RMSE (years) | $R^2$ | CpGs retained |
| --- | --- | --- | --- | --- |
| XGBoost (all 1,000 CpGs) | 4.095 ± 0.285 | 5.501 ± 0.409 | 0.857 ± 0.027 | 1000.0 ± 0.0 |
| Particle swarm optimization | 4.124 ± 0.263 | 5.514 ± 0.364 | 0.857 ± 0.024 | 293.4 ± 24.0 |

**Table 2. Held-out performance over 30 matched train/test splits (mean ± SD).**
| Method | MAE (years) | RMSE (years) | R<sup>2</sup> | CpGs retained |
| --- | --- | --- | --- | --- |
| MOSurvivor | 4.149 ± 0.300 | 5.545 ± 0.392 | 0.855 ± 0.027 | 211.6 ± 54.8 |
| Genetic algorithm | 4.170 ± 0.283 | 5.571 ± 0.389 | 0.854 ± 0.025 | 269.6 ± 45.3 |
| Random search | 4.197 ± 0.313 | 5.611 ± 0.422 | 0.852 ± 0.029 | 278.7 ± 39.2 |

**Figure 1.**
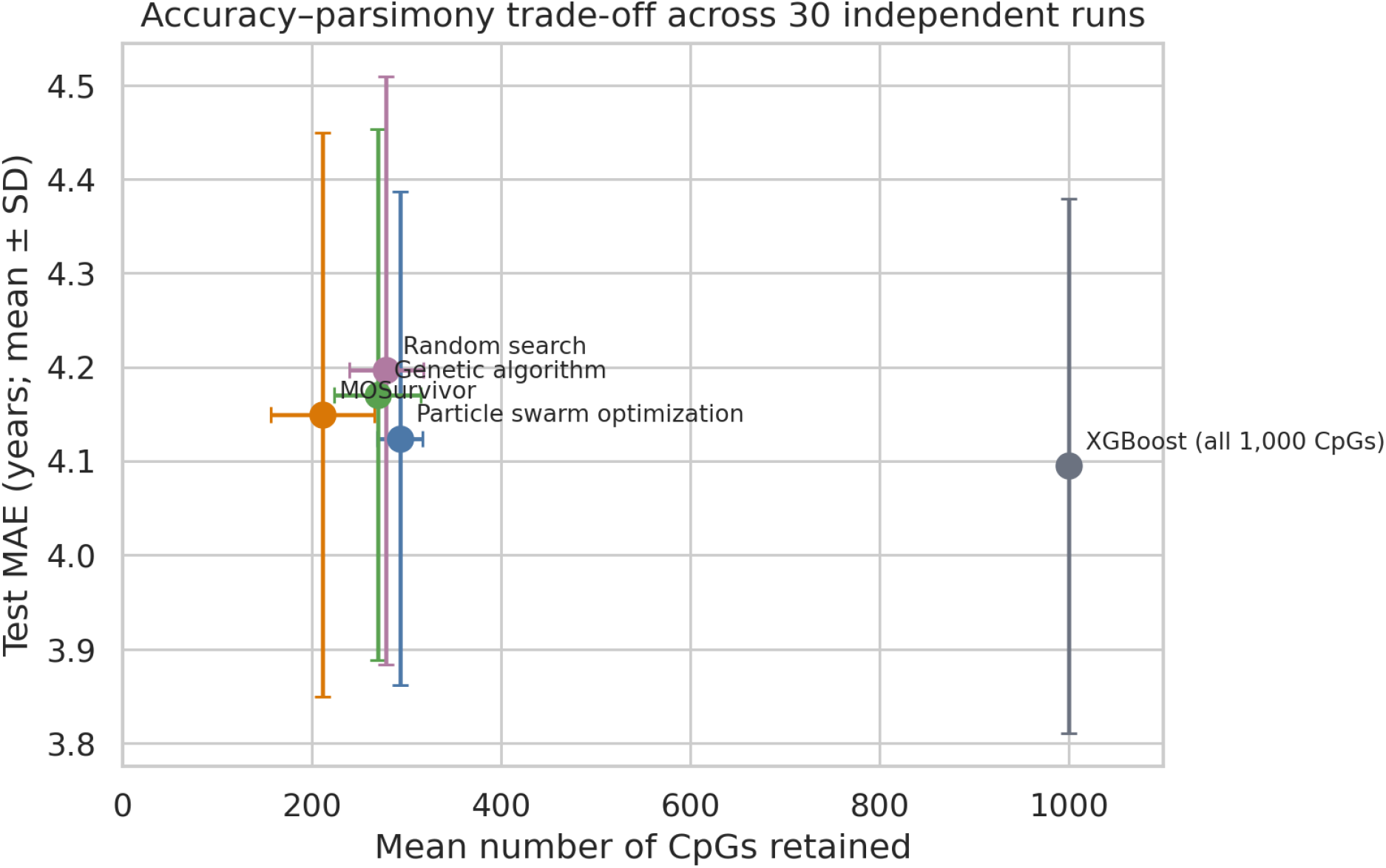
Mean accuracy and subset size. Error bars show one standard deviation across 30 independent runs.

**Figure 2.**
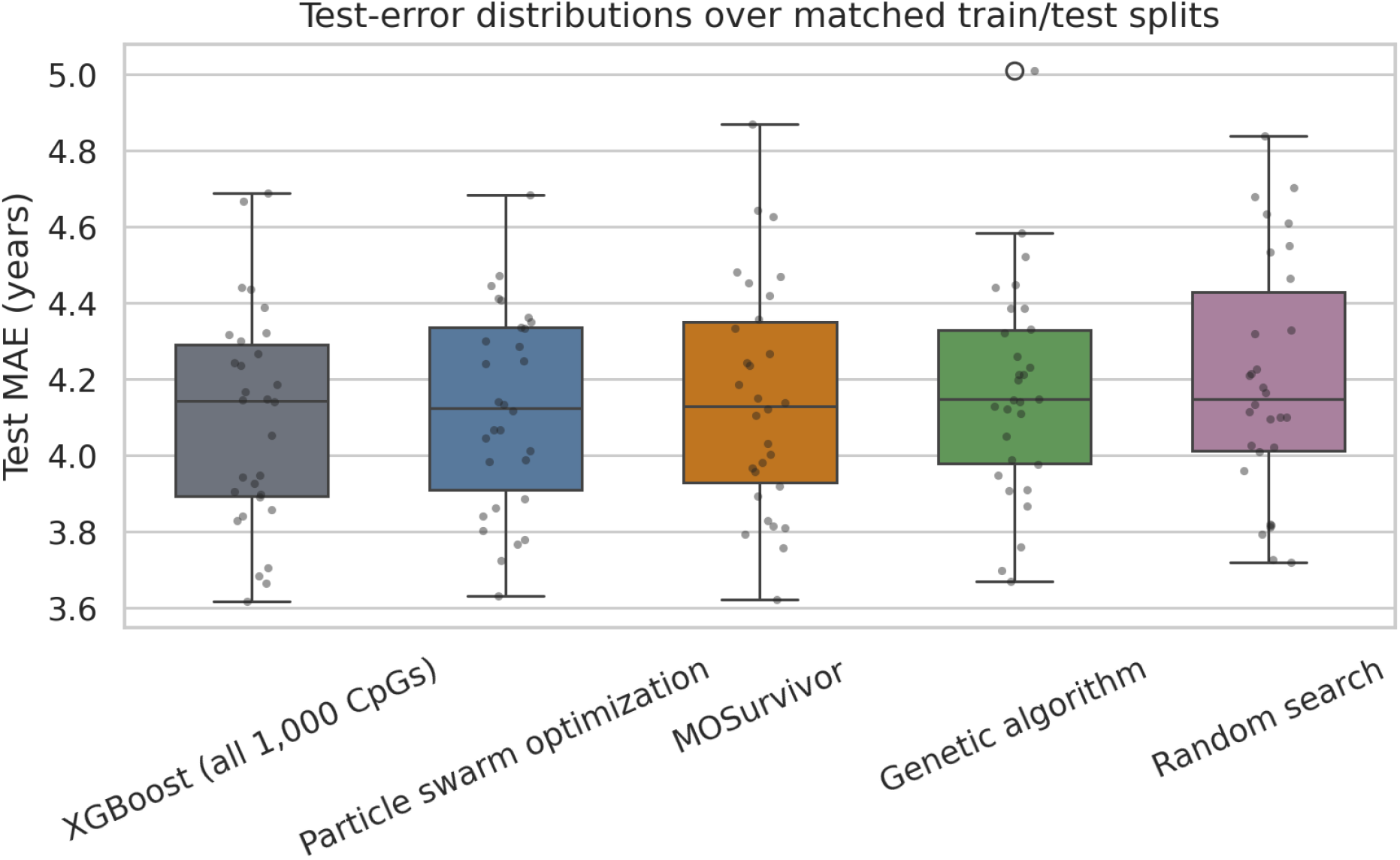
Distribution of held-out MAE values on the same 30 seeded splits.

**Table 3.** Two-sided paired Wilcoxon signed-rank tests across matched splits.

| Comparison | W | Unadjusted p | Holm-adjusted p |
| --- | --- | --- | --- |
| MOSurvivor vs XGBoost (all 1,000 CpGs) | 156.0 | 0.1191 | 0.4763 |
| MOSurvivor vs Genetic algorithm | 200.0 | 0.5158 | 0.9854 |
| MOSurvivor vs Particle swarm optimization | 185.0 | 0.3387 | 0.9854 |
| MOSurvivor vs Random search | 184.0 | 0.3285 | 0.9854 |

### 3.2 Paired statistical comparisons

None of the pairwise differences reached p < 0.05 before or after correction. MOSurvivor produced a lower MAE than full-feature XGBoost on 10 of 30 splits, PSO on 10 of 30, GA on 16 of 30, and random search on 15 of 30. Taken together, the results support accuracy comparability and parsimony, but not a claim of statistically superior prediction.

### 3.3 Feature-selection stability

Across 30 runs, MOSurvivor selected 6,348 CpG instances corresponding to 998 unique loci. Nine loci appeared in at least 15 runs; none appeared in all 30. The highest-frequency locus, cg16867657, appeared in 29 runs, followed by cg04400972 (24) and cg14361627 (20). Despite this recurrent core, the mean pairwise Jaccard similarity among the 435 run pairs was 0.124 ± 0.030 (median 0.123). The low overlap suggests that correlated or partially interchangeable loci yield multiple near-equivalent predictive panels.

**Figure 3.**
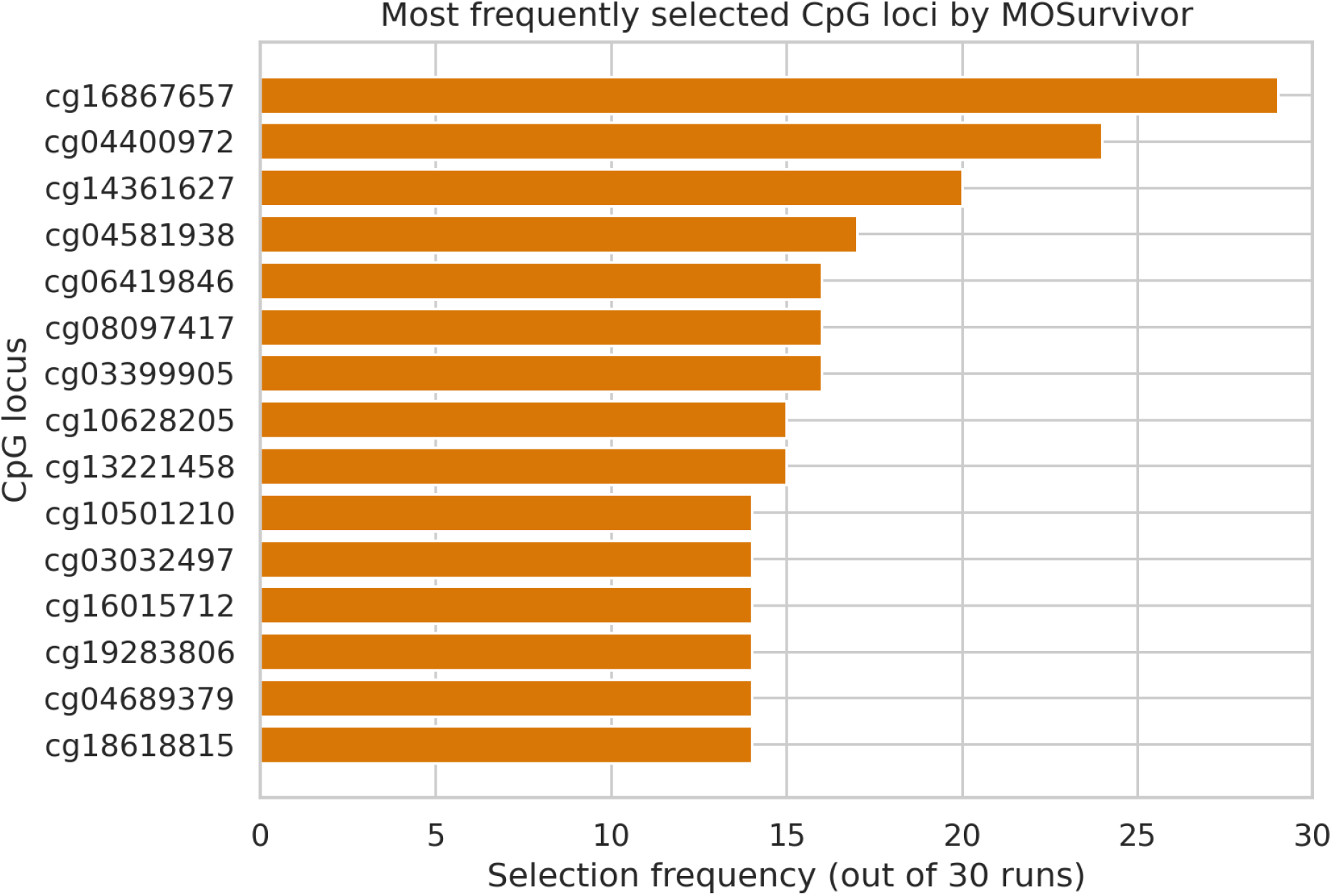
Fifteen CpG loci most frequently selected by MOSurvivor.

## 4. Discussion

The main result is an accuracy-parsimony trade-off: MOSurvivor retained approximately one-fifth of the screened CpGs while preserving nearly the same test error as a model using all 1,000 loci. The 0.054-year difference in mean MAE is small compared with the across-split variability of approximately 0.3 years and was not statistically significant. This is practically relevant because smaller panels can reduce assay design complexity, facilitate interpretation, and narrow the set of loci requiring downstream validation.

MOSurvivor did not produce the best mean predictive accuracy; the full-feature model and PSO were numerically better. The scientific value of the proposed framework lies in joint optimization under parsimony rather than in setting a new accuracy record. That distinction matters because metaheuristic studies can overstate small numerical differences when paired tests are absent. Here, the matched-split analysis and multiplicity correction provide no evidence that MOSurvivor is more accurate than the alternatives.

The stability results reveal two levels of structure. A few loci were repeatedly selected, including cg16867657 in 96.7% of runs, suggesting robust predictive value within this cohort. At the same time, the low average Jaccard similarity indicates high redundancy in the candidate panel. In methylation data, correlated CpGs and age-linked genomic regions can allow distinct subsets to substitute for one another. Frequency alone should not be interpreted as biological causality; annotation to genes and regulatory regions, enrichment analysis, and replication in independent cohorts are required.

Compared with established clocks, the present model is cohort-specific and purely chronological. Horvath’s multi-tissue clock used 353 CpGs across diverse tissues [2], Hannum et al. developed a blood-based predictor from the same general cohort context [1], and very small targeted clocks have also been reported [6]. Direct ranking against these clocks would require applying their published coefficients or validated implementations on exactly the same held-out samples and preprocessing pipeline. The current experiment instead isolates the computational question of whether joint feature and hyperparameter search can compress an XGBoost age predictor.

## 4.1 Limitations and threats to validity

- The initial 1,000-CpG correlation screen used the complete dataset, allowing target information from test samples to influence the candidate panel. Absolute test performance may therefore be optimistic.
- All results derive from one whole-blood cohort and lack external validation across cohorts, laboratories, ancestry groups, age distributions, and array-processing pipelines.
- The 30 runs are repeated random splits of the same 656 individuals, not 30 independent cohorts; uncertainty estimates describe split variability rather than population-level replication.
- Search budgets and wall-clock costs were not available in a form that supports a fair computational-efficiency comparison.
- No biological annotation or pathway enrichment was supplied for the recurrent CpGs, so biological interpretation remains exploratory.
- A fixed scalarization coefficient was used for final archive selection; different clinical or assay-cost priorities could select a different Pareto solution.

### 4.2 Recommended confirmatory protocol

A definitive evaluation should use nested resampling. In every outer training split, CpG correlation screening must be recomputed using training samples only. Inner cross-validation should optimize feature selection and hyperparameters; the outer test partition should remain untouched until final evaluation. At least one independent blood methylation cohort should then be used for locked external validation. Search methods should receive the same number of XGBoost evaluations, and execution time, GPU-hours, calibration, age-stratified error, and subgroup performance should be reported. Finally, recurrent loci should be mapped to genes and regulatory regions and tested for enrichment and replication.

## 5. Conclusion

MOSurvivor jointly optimized an adaptive-threshold CpG subset and XGBoost hyperparameters for epigenetic age prediction. Across 30 matched splits of GSE40279, it retained 211.6 CpGs on average and achieved an MAE of 4.149 years, reducing the 1,000-locus input by 78.8% while remaining statistically indistinguishable in accuracy from the evaluated comparators. Its strongest contribution is model compression with competitive prediction, not superior absolute accuracy. Leakage-free nested selection, equalized search budgets, biological annotation, and independent external validation are required before the framework can support stronger translational claims.

## Declarations

### Data availability

The GSE40279 dataset is publicly available from the NCBI Gene Expression Omnibus at https://www.ncbi.nlm.nih.gov/geo/query/acc.cgi?acc=GSE40279. Run-level results and selected-CpG lists used in this manuscript are available from the corresponding author and should be deposited with the final code release.

